# Imaging of developing human brains with ex vivo PSOCT and dMRI

**DOI:** 10.1101/2024.07.27.605383

**Authors:** Hui Wang, Nathan Blanke, Dayang Gong, Alpen Ortug, Jose Luis Alatorre Warren, Christopher Clickner, William Ammon, Jackson Nolan, Zoe Cotronis, Andre van der Kouwe, Emi Takahashi

**Author notes:** Corresponding authors: Hui Wang, Emi Takahashi, Athinoula A. Martinos Center for Biomedical Research, Department of Radiology, Massachusetts General Hospital and Harvard Medical School, Charlestown, Massachusetts, USA. Department of Radiology, Massachusetts General Hospital and Harvard Medical School, Boston, Massachusetts, USA.

## Abstract

The human brain undergoes substantial developmental changes in the first five years of life. Particularly in the white matter, myelination of axons occurs near birth and continues at a rapid pace during the first 2 to 3 years. Diffusion MRI (dMRI) has revolutionized our understanding of developmental trajectories in white matter. However, the mm-resolution of *in vivo* techniques bears significant limitation in revealing the microstructure of the developing brain. Polarization sensitive optical coherence tomography (PSOCT) is a three-dimensional (3D) optical imaging technique that uses polarized light interferometry to target myelinated fiber tracts with micrometer resolution. Previous studies have shown that PSOCT contributes significantly to the elucidation of myelin content and quantification of fiber orientation in adult human brains. In this study, we utilized the PSOCT technique to study developing brains during the first 5 years of life in combination with ex vivo dMRI. The results showed that the optical properties of PSOCT quantitatively reveal the myelination process in young children. The imaging contrast of the optic axis orientation is a sensitive measure of fiber orientations in largely unmyelinated brains as young as 3-months-old. The micrometer resolution of PSOCT provides substantially enriched information about complex fiber networks and complements submillimeter dMRI. This new optical tool offers great potential to reveal the white matter structures in normal neurodevelopment and developmental disorders in unprecedented detail.

## Introduction

Myelin, which surrounds the axonal fibers, plays a crucial role in brain function by facilitating rapid and coordinated neuronal communication throughout the brain. Postnatal myelination occurs most rapidly within the first two years of life ^1,2^. During the first year, the volume of white matter in the human brain increases by 6–16%, primarily due to myelination ^3^. This time window is critical for maintaining a healthy developmental trajectory of neuroanatomy and physiology ^4^. Disorders of myelination such as multiple sclerosis, Guillain-Barré syndrome, and many others have been associated with a variety of developmental and cognitive impairments ^3^. MRI has revolutionized the way myelination is assessed in young children. Prior to the use of MRI, the only possible way to determine the developmental status of myelination was by clinical course, neurological examination, and histological examination ^5^. Diffusion MRI (dMRI) sequences use diffusion-sensitized gradient pulses to probe the anisotropic diffusion of water molecules in brain tissue, which allows imaging of pathways with coherent water diffusivity, including developing axons. Fractional anisotropy (FA), which indicates the degree of anisotropy of water diffusivity, serves as a good indicator for white matter organization, because coherent axonal organization restricts the movement of water molecules perpendicular to the axon bundle ^5^. A link between FA and myelin maturation in infants has been suggested ^6–8^. However, rapid changes in brain anatomy and physiology pose many distinct limitations to the acquisition, data processing, analysis, and interpretation of these developmental trajectories ^4^. It is known that dMRI tractography has difficulty identifying complex fiber configurations from multiple directions in complex structures^9^. This problem becomes more prominent in the developing small brain.

Optical imaging methods provide superior resolution compared to MRI techniques. Among them, polarization sensitive optical coherence tomography (PSOCT) provides label-free and depth-resolved imaging contrasts that originate from light scattering and tissue birefringence ^10^. When applied to brain imaging, the intensity contrast from PSOCT reveals the gross anatomical structure and provides quantification of the scattering coefficient ^11,12^. Birefringence originates from the structural anisotropy of myelinated fibers and results in an optic axis that is parallel to the fiber orientation ^13^. Fiber tracts as small as tens of micrometers in diameter can be resolved with the retardance map, and the fiber orientation can be measured by the optic axis orientation ^14,15^. The development of automatic serial sectioning PSOCT (as-PSOCT) based on blockface imaging has proven to be a valuable tool for mapping the complex and intricate white-matter organization in the human brain ^16^ (also see a recent review ^17^). By integrating a tissue slicer into the imaging system, as-PSOCT has proven to be an effective way to map fiber orientations across cubic centimeter specimens of human brain. One advantage of as-PSOCT is that it does not suffer from the nonlinear distortions plaguing slice-based histological techniques that demand complex registration frameworks to correct ^18,19^. As a result, the as-PSOCT technique allows for the reconstruction of large volumes of brain samples with microscopic-level resolution. To date, PSOCT has been used to study adult human brains. The usefulness of PSOCT in infancy and early childhood, when myelination is in process, has not been reported.

In this work, we utilized as-PSOCT, in combination with dMRI, to investigate the neurodevelopment of the human brain in early childhood. We found that as-PSOCT was sensitive in delineating the myelination process in the infant brain as early as 3 months old. Optical properties revealed distinctive developmental patterns in different brain regions across the first 5 years. Correlations between as-PSOCT and dMRI images showed general agreement in fiber tracts, while the optical imaging resolved finer details on tissue properties and fiber orientations. The high resolution of the as-PSOCT technique offers great potential for studying typical and atypical neurodevelopment in the human brain.

## Method

### 2.1 Samples

Five human brain samples (3 males, 2 females) were obtained in coronal slabs from the University of Maryland Brain and Tissue Bank (UMBTB) through the NIH NeuroBioBank (NBB) network. The slabs were approximately 2 cm thick and fixed in 10% formalin. The brain samples had different developmental timelines: 3 months old, 6 months old, 15 months old, 50 months old, and 54 months old. The latter two were considered age matched. One subject was diagnosed with sudden infant death syndrome (3 months old), one with autism spectrum disorder (54 months old), and the other three were controls without neurological disease. The postmortem intervals were less than 40 hours. The samples were submerged in fomblin for the dMRI scan. After dMRI, one coronal slab per brain was embedded in agarose using a customized grater ^20^ and refractive index matched in either 30% TDE or 50% glycerol for PSOCT imaging. Supplementary Table 1 describes the detailed individual information about the specimens.

### 2.2 Ex vivo diffusion MRI and data processing

Slabs of the postmortem brains were prepared for MRI scanning by soaking in bags containing fomblin oil (same used as, e.g., ^21,22^). These bags were put in a container for each subject, arranged side by side with plastic plates dividing them. The scans were performed at the Athinoula A. Martinos Center for Biomedical Imaging for all 5 brain samples. The primary obstacle to conducting high-resolution ex vivo human dMRI studies is the significantly reduced T2 and diffusivity of fixed tissue ^23^. To address these challenges, we adjusted the acquisition parameters by referring to the previous literature on fixed human brain dMRI, and further optimized them based on our previous study ^22^. Diffusion-weighted MRI data were acquired using a diffusion-weighted SSFP sequence ^24,25^ on a 3 Tesla Siemens Trio scanner. The following parameters were used: TR = 28.82 ms, TE = 24.42 ms, flip angle = 35°, resolution = 0.8 mm isotropic, in-plane FOV = 144 mm × 144 mm, number of slices = 88, bandwidth = 149 Hz/px. Diffusion weighting was performed along 60 directions with 10 T2-weighted b = 0 measurements. In diffusion-weighted SSFP acquisitions, the b-value varies with tissue T1 and T2 relaxation times. In our experiment, with diffusion gradient duration of 20 ms, the b-value was approximately 4,000 s/mm^2^. Total scan time was 6h 11min 43s.

Diffusion Toolkit ^26^ and TrackVis (Version 0.6.1; trackvis.org) were used to reconstruct and visualize tractography. The FACT algorithm and 60°-angle thresholds were used in the diffusion tensor imaging (DTI) model to reconstruct tractography pathways. No FA threshold was applied ^27^. As multiple brain slabs were scanned together in a box, after tractography was performed in the entire FOV, individual slabs were segmented using Amira software ^28^ (3D Version 2021.2, Thermo Fisher Scientific, Waltham, MA, USA), and were analyzed separately.

### 2.3 PSOCT imaging and data processing

#### PSOCT system

One slab per sample was imaged with PSOCT. The anatomical locations of the coronal slabs scanned using both modalities were labeled in Fig 1. A home-made automatic serial sectioning PSOCT (as-PSOCT) system was used for data collection. The system integrates a commercial spectral domain PSOCT system (TEL220PS, Thorlabs), motorized xyz translational stages, and a vibratome tissue slicer. Custom-built software, written in C++, provides coordinated data acquisition, xyz-stage translation, and vibratome sectioning for automatic imaging of brain blocks. The maximum sensitivity of the PSOCT was 109 dB. The imaging depth was 2.6 mm with an axial resolution of 4.2 μm in tissue. The samples were imaged with a scan lens objective (OCT-LSM3, Thorlabs), yielding a lateral resolution of 10 μm. One volumetric acquisition was composed of 350 A-lines and 350 B-lines covering a field of view (FOV) of 3.5 × 3.5 mm at an A-line rate of 50 kHz. We imaged the block face surface of the entire coronal section via tile scans with a 20% overlap. Two of the five samples (50 months and 54 months old) went through serial sectioning to acquire large volumetric image sets. A 100–150 µm thick slice was removed from the tissue surface by the vibratome to expose the deeper region until the whole block of tissue was imaged.

**Fig. 1.**
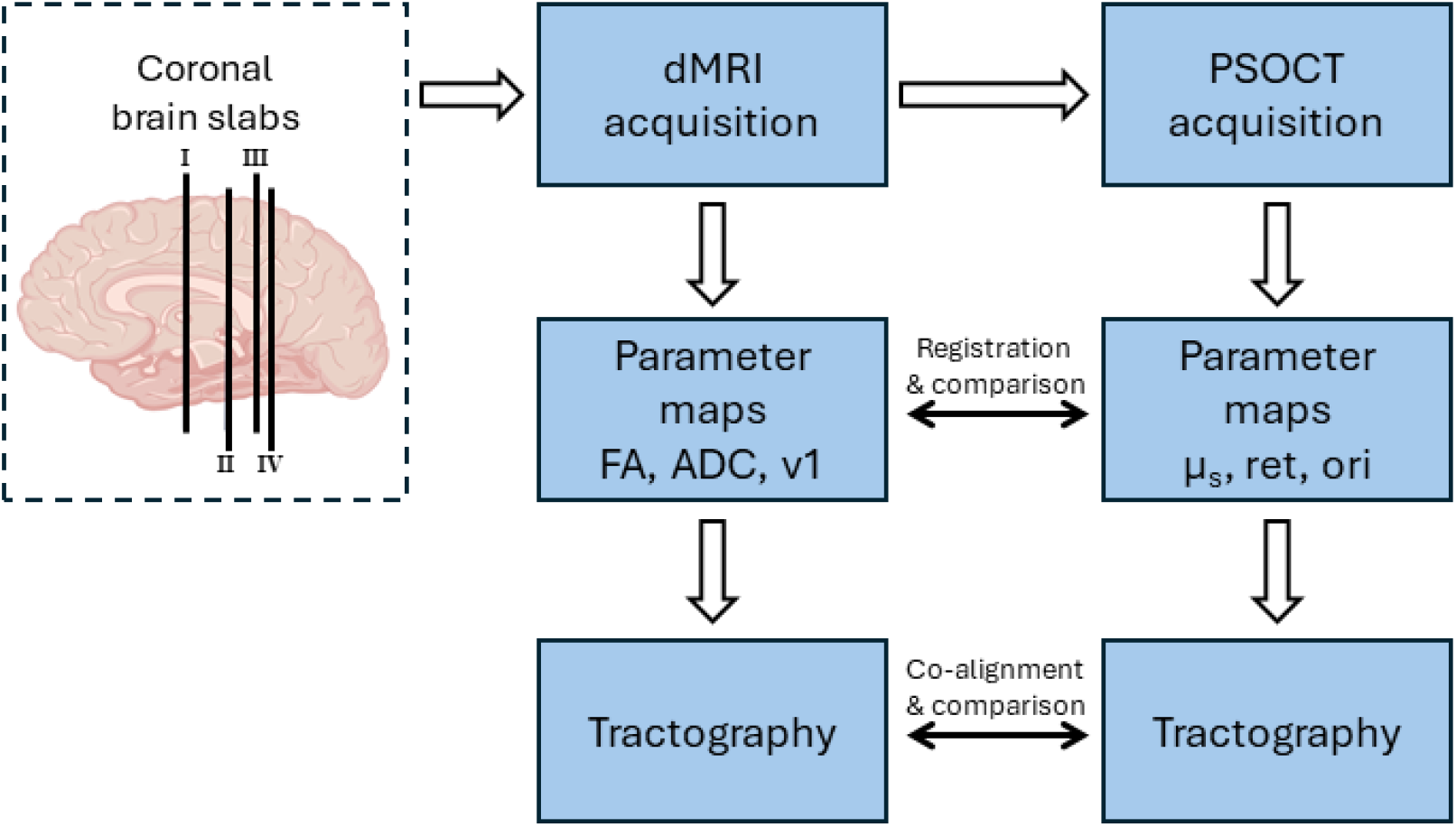
Pipeline of image acquisition and processing. The anatomical location of each coronal slab is overlaid on the cartoon diagram of the brain.

#### Image reconstruction

Inverse Fourier transform of interference-related spectral oscillations yielded complex depth profiles in the form of *A*_1,2_(*z*) exp[*iϕ*_1,2_(*z*)], where *A* and *ϕ* denote the amplitude and phase as a function of depth *z*, and the subscripts represent the polarization channels. The image contrasts of intensity A(*z*), *R*(*z*), retardance, *δ*(*z*), and optic axis orientation, *θ*(*z*) along depth, were obtained by *R*(*z*) ∝ *A*_1_(*z*)^2^ + *A*_2_(*z*)^2^, *δ*(*z*) = arctan[*A*_1_(*z*)/*A*_2_(*z*)] and *θ*(*z*) = [*ϕ*_1_(*z*) − *ϕ*_2_(*z*)]/2, respectively. We also quantified the voxel-wise attenuation coefficient following the method of Vermeer et al. In the near-infrared spectral range, light attenuation within the tissue is dominated by scattering, whereas absorption is negligible. Therefore, we used the scattering coefficient (μ_*s*_(*z*)), calculated as 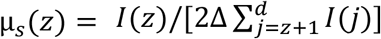, to represent the attenuation coefficient, where *z* is the pixel number in depth, *I* is the reflectivity signal, Δ is the pixel size in depth, and *d* is the total imaging depth in mm ^29^. En-face scattering coefficient and retardance images were calculated by averaging the respective values along the slice thickness. The orientation at each pixel in the en-face axis orientation images corresponded to the peak of a histogram constructed by binning the measured orientation values into 5° intervals. We stitched the tiles to reconstruct the entire section in the Fiji software which computed the overlap between tiles and used linear blending for image fusion ^30^. In the two samples with serial sectioning, the images of individual slices were stacked together to render the volumetric reconstruction of the coronal slab.

#### Tractography

Tractography was applied to the in-plane optic axis orientation images using the conventional method for DTI modelling in Diffusion Toolkit. The optic axis orientation was used to track the fibers, while the orientation for the z-direction was set to 0. The fiber tracking algorithm is based on the spherical harmonic basis method ^31^. Tracts were created with a maximum angular threshold of 45° for tracking and masked by the retardance image to include the white matter only.

### 2.4 PSOCT-dMRI registration

We co-registered PSOCT and dMRI data for a further correlation and comparison study. For samples that only had one PSOCT imaging section, we found the closest dMRI slice to the PSOCT images and manually rotate the parameter images and the tractography to match the anatomical directions. For samples that had PSOCT with serial sectioning, we used robust registration, an automatic registration method that is insensitive to outlier areas of the images ^32^, to register volumetric PSOCT and dMRI. We aligned the dMRI and PSOCT data from each block by registering the apparent diffusion coefficient (ADC) map to the scattering coefficient map, as they possess the best gray/white matter contrast in the younger ages. The absence of non-linear distortions is a key feature of the PSOCT acquisition that facilitates this step, in comparison to alternative histological techniques. Thus, an affine registration was sufficient for the cross-modal alignment ^33^. To evaluate the volumetric registration, we created a tissue mask and white matter mask for both PSOCT and dMRI volumes and computed the Dice coefficient ^34^ between the two modalities. For the dMRI orientation registration, we applied the rotational component of the affine transformation to the dMRI orientation vectors and extracted the corresponding in-plane orientation. This allowed voxel-wise correlation of the diffusion-based orientation with direct measurements of axonal orientations from PSOCT. We also compared the PSOCT orientation with dMRI tractography.

### 2.5 Correlation between PSOCT and dMRI

To quantitatively evaluate the correlation of the metrics between PSOCT and dMRI, we conducted two analyses. We manually segmented the white matter in both PSOCT and the corresponding dMRI slice. For the samples that had volumetric co-registration between the two modalities, we also manually selected 0.9 × 0.9 mm ROIs that were evenly distributed in the white matter in corresponding voxels of PSOCT and dMRI images. For PSOCT, we calculated the mean scattering coefficient, retardance and circular mean ^35^ optic-axis orientation of each ROI. For dMRI, we calculated the mean ADC, FA and circular mean diffusion orientation of each ROI. It is noted that the dMRI orientation vector was mapped into the PSOCT imaging plane first before obtaining the circular mean diffusion orientation. We then correlated the optical properties and dMRI parameter metrics using a linear fitting tool, both within the sample and across samples of different ages. We report the residual sum of squares and Pearson’s r values as metrics of linear fitting quality. For quantitative comparison of orientation, we investigated the angular difference between the two modalities and displayed them in a polar plot. The full pipeline of data acquisition and image processing is summarized in Figure 1.

## Results

### 3.1 PSOCT showing myelin development in the first 5 years

The optical properties of PSOCT reveal detailed structural components in the brain. It is known that both scattering coefficient and birefringence are high in myelinated fibers and that retardance provides a measure of the degree of alignment in myelinated fibers across the imaging depth. In this study, we show that the optical properties of PSOCT reveal the myelination process in young children. The scattering coefficient in white matter increases with age (Fig 2, top row). At 3 months of age, the contrast between white and gray matter is not outstanding (a), whereas by 6 months of age, white matter starts to show a higher value than the gray matter in some brain regions (b). It is interesting to observe the regional heterogeneity of myelination during infancy. In the anterior part of the brain (section I), the border of the parietal/frontal lobes (somatosensory/motor cortices) show a higher scattering coefficient than the temporal lobe, indicating that myelination is advanced in the superior part of the anterior section at 6 months of age. This regional difference becomes invisible in the posterior brain by 15 months of age (c). The scattering coefficient in the white matter continues to rise substantially in the next four years (d and e). In contrast, the scattering coefficient stays low in the gray matter despite a small increase across multiple years. Quantitative analysis shows that the value increases almost 10 times in the white matter across the first five years of life (Fig 3a). The retardance maps present more varying contrasts both within and across subjects, with white matter values higher or lower than gray matter (Fig 2, bottom row). One possible contributing factor is the heterogeneity of optic axis orientation within the fiber bundles across the imaging depth. In regions where multiple fiber tracts meet and where fibers are oriented through the imaging plane, a reduced retardance is observed due to signal cancellation at the oppositely oriented myelin optic axes ^36,37^.

**Fig. 2.**
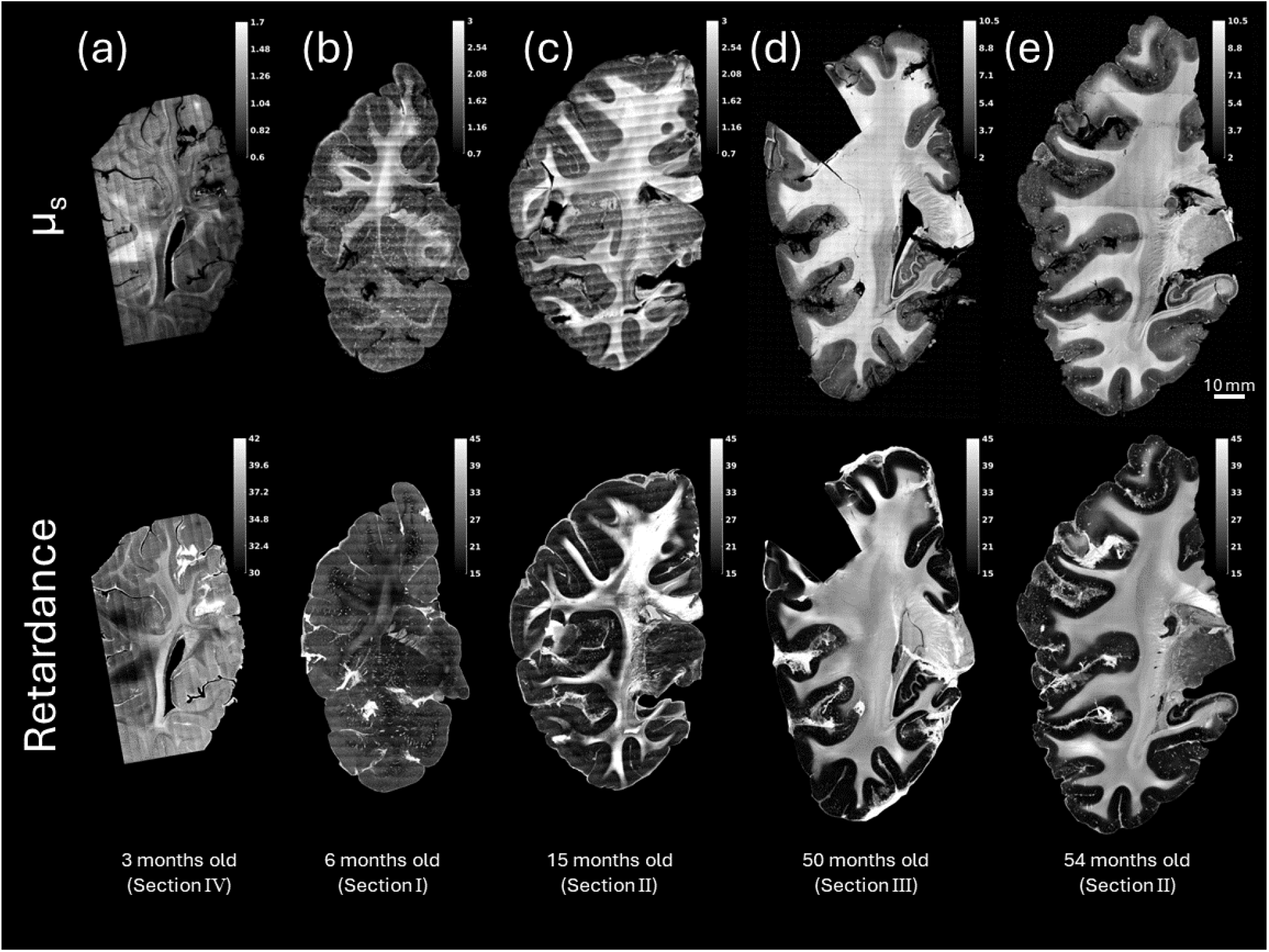
PSOCT optical property maps of scattering coefficient and retardance showing brain development in the first 5 years. (top) Scattering (μ_s_) maps and (bottom) retardance maps are shown for five samples from infancy to early childhood: (a) 3 months old, (b) 6 months old, (c) 15 months old, (d) 50 months old, (e) 54 months old. The scale for each grayscale image is shown in the top-right inset of each panel. The anatomical location of each slab is denoted by Roman numerals and is shown in Fig. 1.

**Fig 3.**
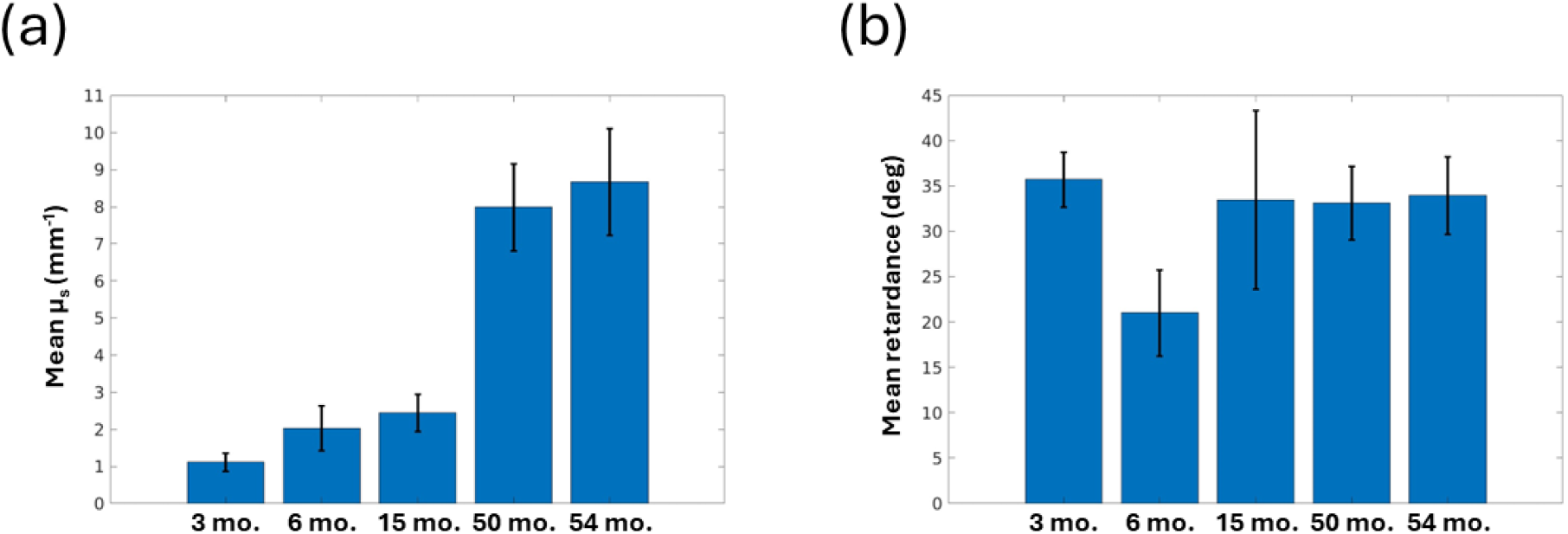
Change of PSOCT optical properties with white matter development. (a) Mean scattering coefficient (μ_s_) and (b) mean retardance analyzed across all pixels in white matter. The error bars show standard deviation.

### 3.2 Macroscopic and microscopic fiber pathways imaged with PSOCT and dMRI

The myelin sheath around axons presents an optic axis that is normal to the fiber orientation ^38,39^, and that anisotropic material property provides the ability to map the in-plane fiber orientation with PSOCT optic axis orientation maps. The optic axis orientation of fiber pathways has been well-studied in adult human brain with PSOCT and other polarization microscopy techniques ^36,37,40^. Here we show that the optic axis enables fiber orientation measurements during the first 5 years of life (Fig 4, top row). As indicated by the color wheel, the major fiber bundle orientations seen in multiple samples are consistent across different ages. Large fiber bundles of the internal capsule running along the superior-inferior axis show red-magenta colors, while tracts extending left and right into different cortical regions show colors of green, blue, and yellow. It is noted that the low myelinated white mater regions show greater noise in the orientation measurement. The optic axis orientation in the 3-month-old (Fig. 4a, top) only manifests in major fiber bundles that share orientation in the plane, and the overall noise level is higher than in the rest of the samples. At 6 months of age (Fig. 4b, top) the SNR in the temporal lobe is lower than in the parietal lobe, likely due to different myelin levels. Despite the differences in noise levels, our PSOCT technology proved sensitive in capturing fiber orientation in infants as early as 3 months of age.

**Fig 4.**
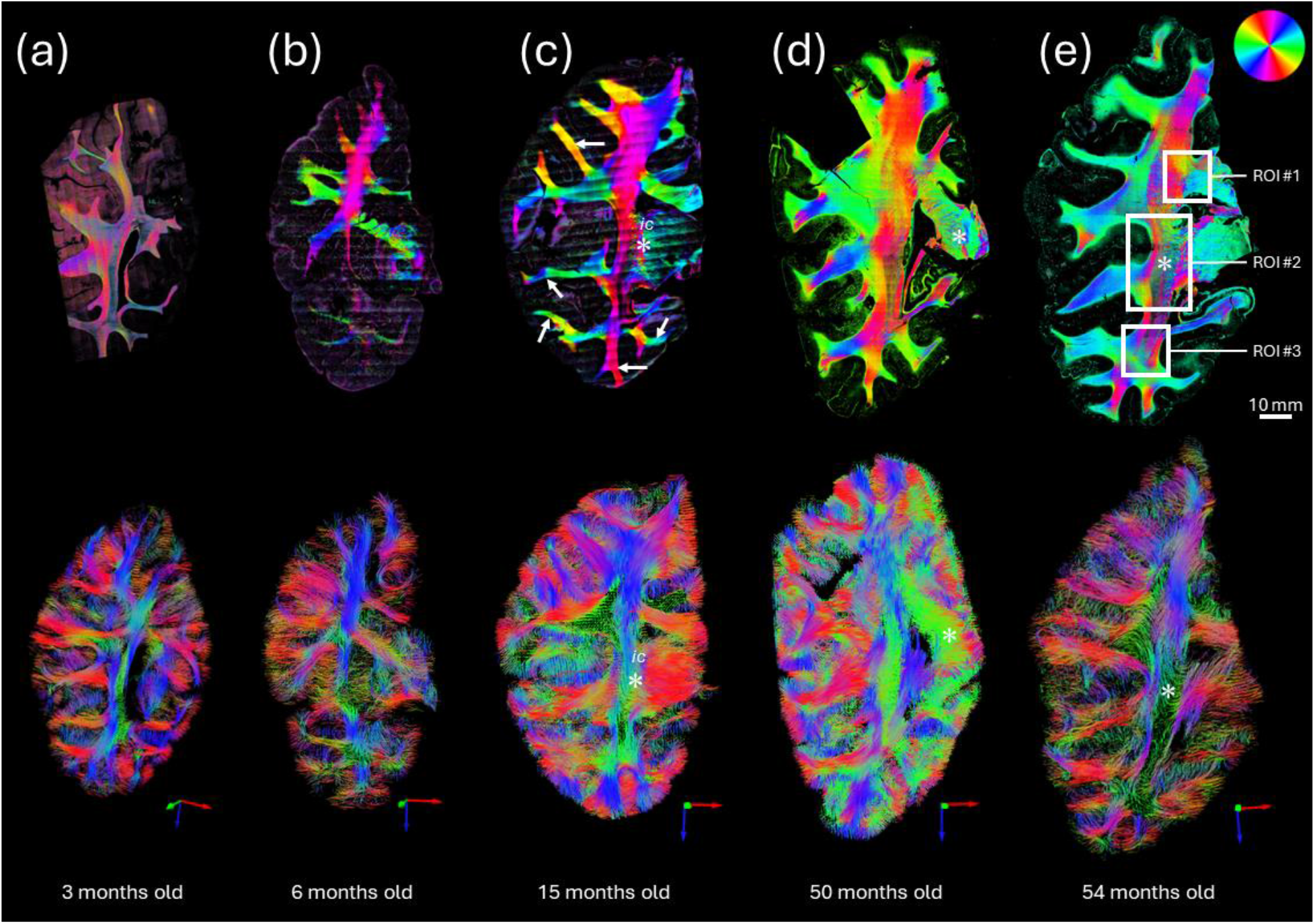
Fiber pathways visualized with PSOCT optic axis orientation maps and dMRI tractography. (top) PSOCT optic axis orientation maps and (bottom) dMRI tractography visualizations are shown for five samples from infancy to early childhood samples: (a) 3 months old, (b) 6 months old, (c) 15 months old, (d) 50 months old, (e) 54 months old. The color scale used for showing the fiber orientation based on PSOCT optic axis is shown in the top-right corner of (e). The color axis used to display each dMRI tractography view is shown in the bottom-right of each inset. Three regions of interest are indicated in (e) that were selected for closer inspection in Fig. 5.

In each sample, the dMRI tractography was aligned with the PSOCT imaging plane for comparison. Overall, the dMRI-based tractography was consistent with the PSOCT-based orientation. For example, the main axis of the internal capsule (ic) is superior-inferior (Fig. 4), and fibers toward the outer gray matter are oriented either left-right or superior-inferior (white arrows), depending on location. dMRI maintains a high quality of tract identification in the younger samples at 3 months old (Fig. 4a, bottom row) and 6 months old (Fig. 4b, bottom row). In contrast, PSOCT detected more detailed and distinct fiber orientation in samples at 15 months old and older (Fig. 4c-e, asterisks).

The high resolution of PSOCT allows us to examine detailed fiber configurations in the developing brain. Figure 5 shows the fiber orientation maps of the 54-month-old in three anatomical regions along the medial side of the coronal slab, from superior to inferior, including (1) the junction of the corpus callosum and projection pathways, (2) the peri-internal capsule, and (3) the temporal stem pathways at the temporal horn of the lateral ventricle junction. Region 1 (Fig. 5, top) has three groups of large fiber bundles all coming from different directions and meeting in the center of the ROI. These three fiber bundles have clear but non-homogeneous boundaries in the orientation color map, which predicts that they intersect each other and change their layout in depth at the intersection. Similarly at the intersection (d, white arrow), dMRI tractography is blank (Fig. 5d, top), possibly due to a high degree of crossing. Although our current PSOCT measurements are limited to the in-plane orientation, the spatial pattern of the orientation map provides useful insights about the fiber configurations in regions where multiple fiber groups meet together. In Region 2, which covers the internal capsule and surrounding regions, multiple bands of fiber groups are elaborated (illustrated by the dotted curves), organized along the left-right axis and presented by altered orientation structures and SNR. On the left side, there are two thin layers of coherent fiber tracts with different orientations [red and green in PSOCT tractography (Fig. 5c, middle)]. In the right half, there are small tracts crossing with each other at different orientations. There is a significant SNR drop in the middle of the fiber tracts where scattering coefficient map (Fig. 5a, middle) shows a dim intensity. The same region exhibits a lack of dense fiber tracts in the PSOCT tractography (Fig. 5c, middle), suggesting that fiber tracts running through the plane are missing. There is another group of fiber tracts running mostly horizontally. The rightmost band is composed of highly interwoven fibers running through the thalamus region. The dMRI tractography captures the main fiber orientations but misses the delicate crossing patterns (Fig. 5d, middle). Region 3 (Fig. 5, bottom) is located where most of the fibers are oriented similarly. Despite this similarity, we still observe different fiber groups in the PSOCT tractography (Fig. 5c, bottom), as they present altered SNR as shown in the axis orientation (Fig. 5b, bottom). Low intensity on the scattering coefficient map (Fig. 5a, bottom) indicates regions where different fiber configurations are present but are not captured with the PSOCT tractography. dMRI tractography indeed identifies tracts running through the plane in this region (Fig. 5d, bottom).

**Fig 5.**
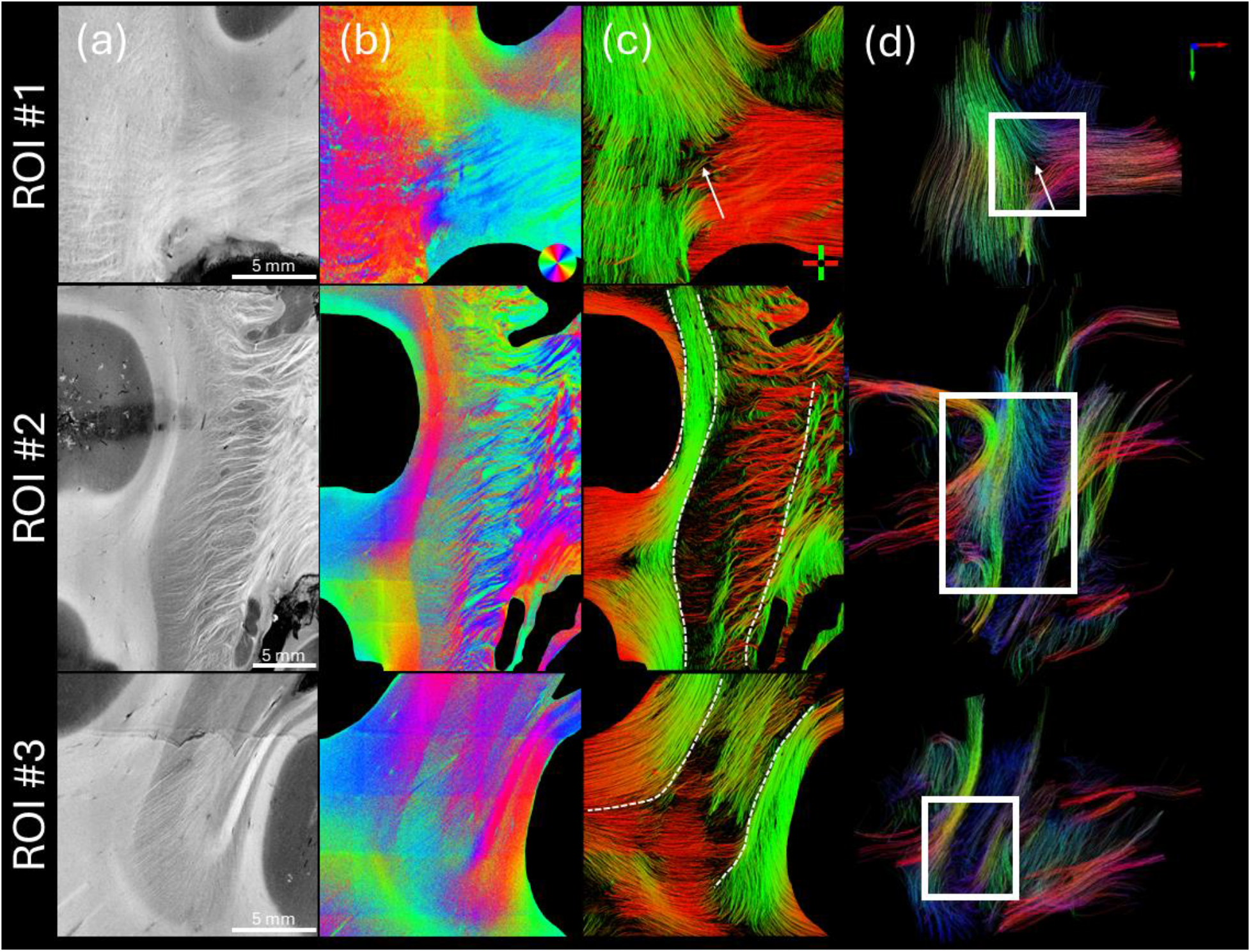
Microscopic fiber orientation and tractography. In three regions of interest (indicated in Fig. 4e), images are shown for (a) PSOCT scattering coefficient map, (b) PSOCT optic axis orientation, (c) PSOCT tractography, and (d) dMRI tractography. The color map used to show in-plane PSOCT fiber orientation is shown in the bottom-right corner of (b). The color axis used to display the PSOCT tractography is shown in the bottom-right corner of (c), and the one used to display the dMRI tractography is shown in the upper-right corner of (d).

### 3.3 Quantitative correlation between PSOCT and dMRI

As the results in section 3.1 show, the optical property of scattering coefficient increases with age. Previous dMRI studies have suggested that ADC values decrease, and FA values increase during postnatal development. Therefore, we are interested in examining the correlation between optical properties and dMRI parameter maps over the first few years of life. We analyzed the scattering coefficient and the retardance in the white matter and the ADC and FA values from the corresponding slice. A scatter plot shows a strong negative correlation between the optical properties and the ADC value (Fig. 6a, c), and a moderate positive correlation between the optical properties and the FA value (Fig. 6b, d).

**Fig 6.**
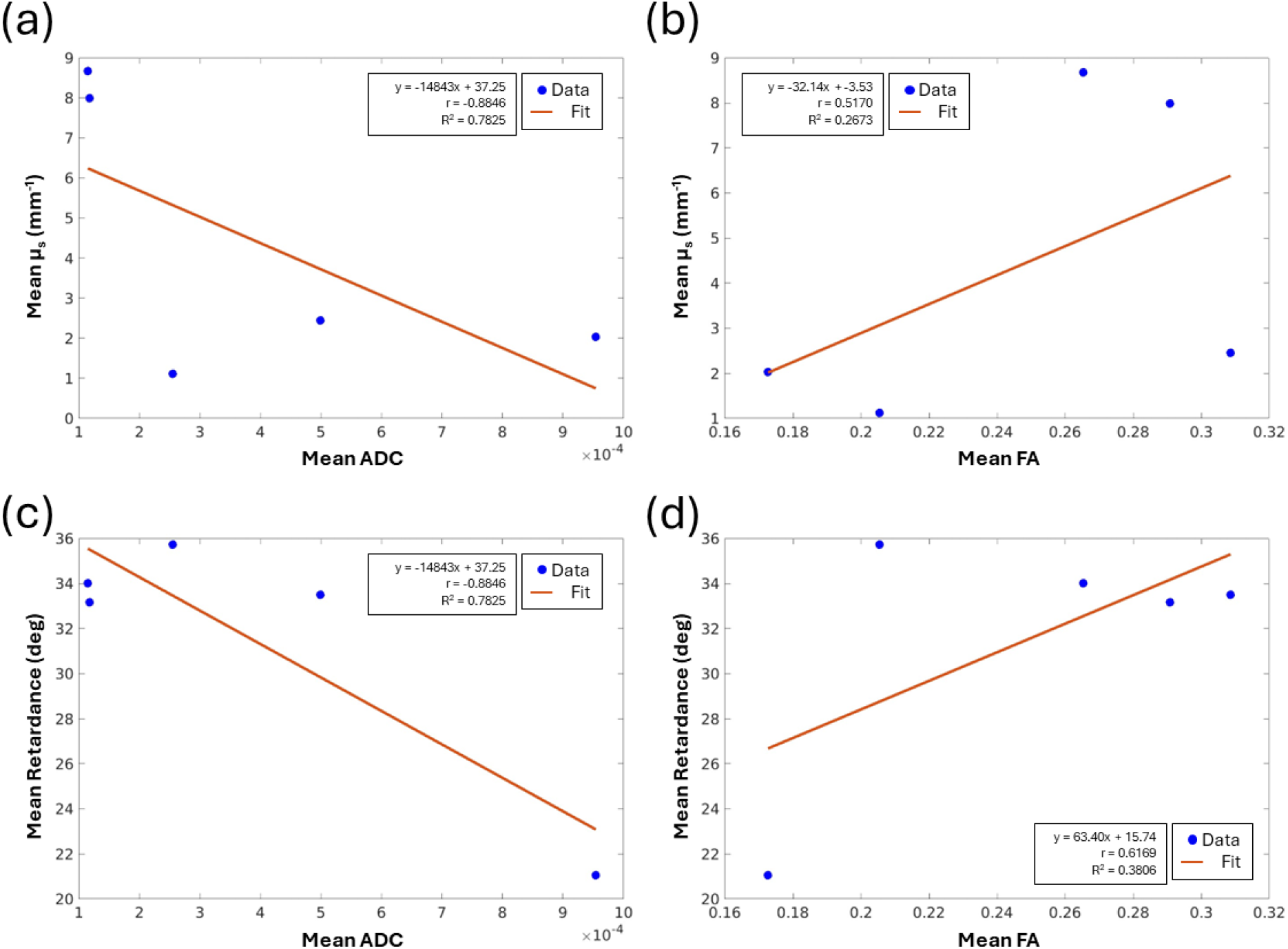
Correlation between optical properties (µ_s_ and retardance) and dMRI parameters (ADC and FA) across the 5 samples of different ages. Blue dots are scatter plot and red line is the linear fit of the scatter plot.

We also investigated the within-subject correlation of the parameter maps as a means of examining the regional variability addressed with the two imaging modalities. To make a quantitative comparison, we first assessed the quality of PSOCT and dMRI co-registration in the two volumetric images, using the Dice coefficient on both the tissue masks and the white matter masks. The Dice coefficient on tissue mask was 0.96, indicating an almost perfect overlap of the co-registered datasets. The Dice coefficient of the white matter masks was 0.87. Figure 7 demonstrates the co-registration results for one section at 54 months of age.

**Fig 7.**
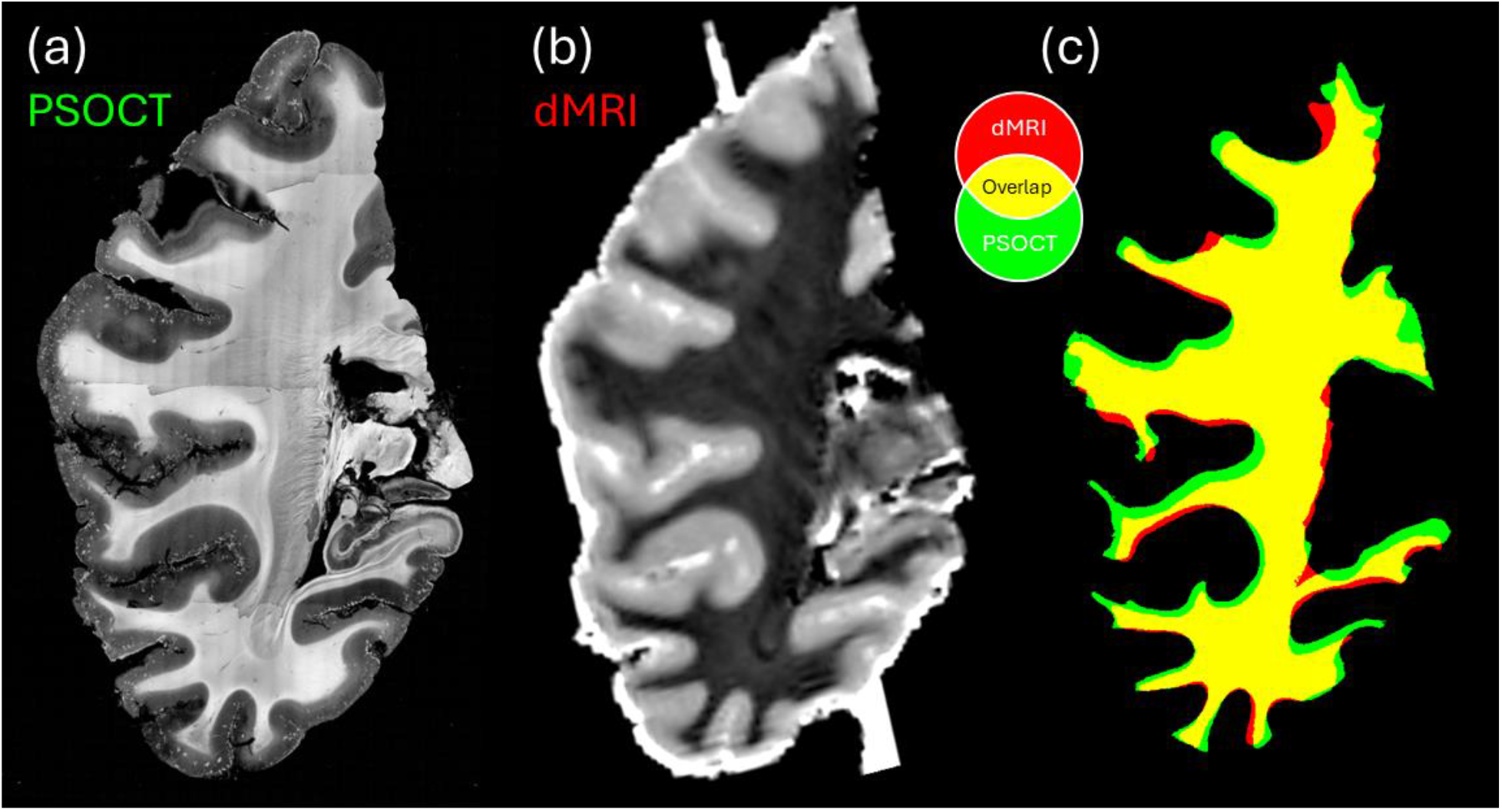
Registration between PSOCT and dMRI images. Registration between (a) PSOCT and (b) dMRI was performed based on their tissue and (c) white matter masks. (c) The overlap between the PSOCT and dMRI white matter masks following registration.

Next, we selected 120 ROIs dispersed in the white matter of each sample and examined the relationship between PSOCT optical properties and dMRI parameter maps, including retardance vs. FA, retardance vs. ADC, scattering coefficient vs. FA, scattering coefficient vs. ADC, and PSOCT orientation vs. dMRI orientation. Among the scalar maps we examined, only retardance and FA show a moderate correlation in the white matter regions (Fig. 8a, c). These two metrics may reveal the degree of structural anisotropy that is higher in coherent fiber bundles and lower in crossing regions. Although weakly correlated with other scalar metrics, the fiber orientation shows a strong agreement in regional fiber bundles between the two modalities. To compare the distribution of in-plane fiber orientation angles between PSOCT and dMRI, we plotted the angular difference between the two modalities (Fig. 8b, d), which minimizes the difference in polar space for accurate representation (i.e., 178° – 1° = 3°, not 177°). The angular differences between dMRI and PSOCT are distributed in a narrow range centered at 0°. The consistency in the measurements of fiber orientation serves as a cross-validation for the two modalities to study the connective pathways in the developing brain.

**Fig 8.**
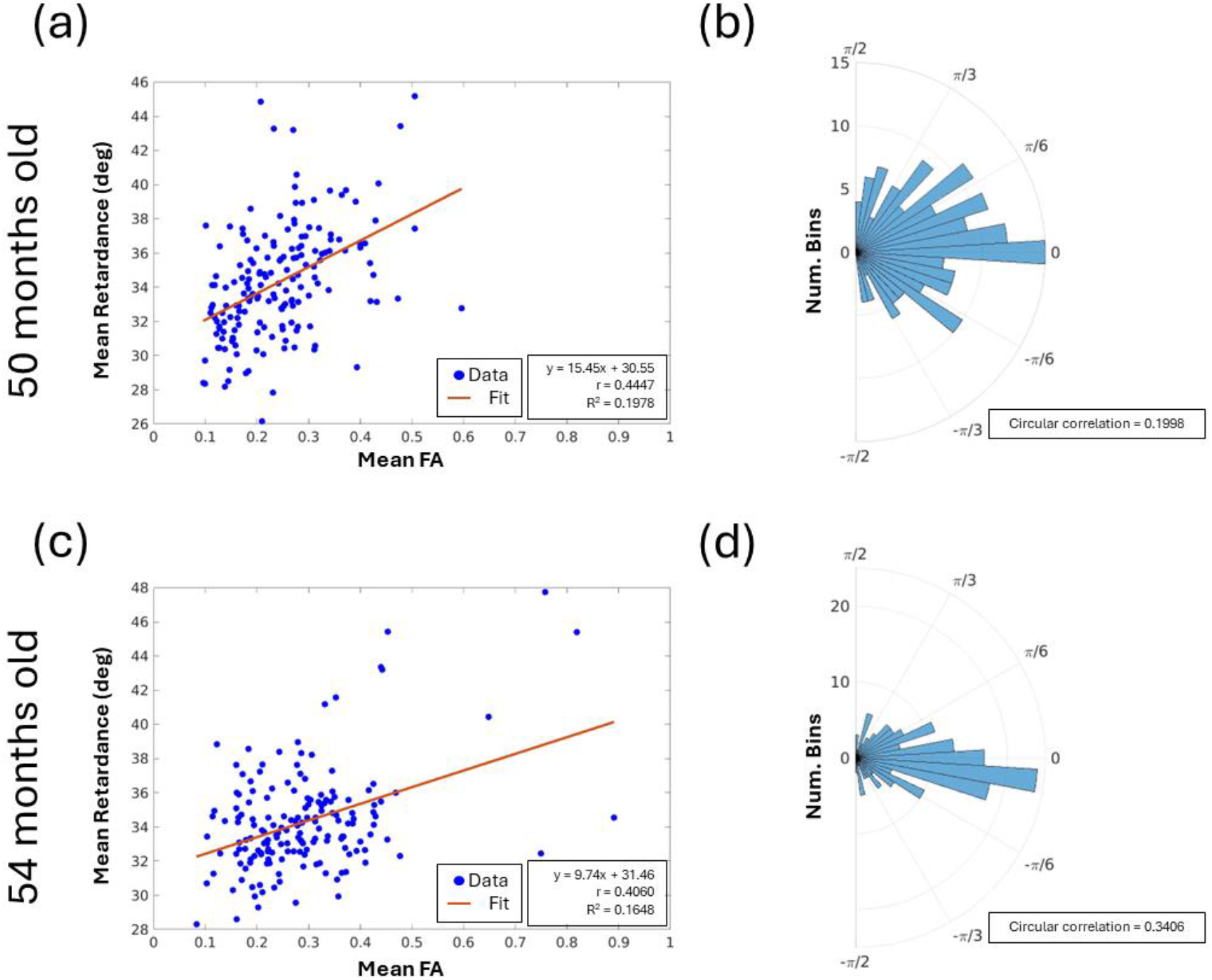
Comparison between PSOCT and dMRI parameter maps. PSOCT and dMRI images were registered for the (top) 50-month-old and (bottom) 54-month-old samples and compared in white matter. (a,c) Mean PSOCT retardance vs. mean dMRI FA. (b,d) Angular difference between PSOCT orientation and dMRI orientation.

## Discussion

The use of PSOCT to reveal microstructures and fiber pathways in large-scale brain tissues has received considerable attention in human neuroscience recently. The advantageous blockface imaging and label-free contrast mechanism enable histological level details in the image, while remaining free from drawbacks like staining bias and limiting tissue distortions or damage encountered in traditional histology. In this study, we leveraged the PSOCT technology to investigate the neurodevelopment of white matter during the first 5 years of life. We used a combination of PSOCT and dMRI to study five whole coronal slabs of the human brain across different ages. PSOCT clearly advocates for the use of optical properties as important biomarkers for myelination processes. The optic axis orientation delineates the direction of the fiber tracts in the infant brain at the age of 3 months, which is consistent with dMRI. The microscopic resolution of PSOCT is capable of resolving small fascicles less than 100 micrometers thick, as well as complex fiber configurations with inter-weaving and splitting. To the best of our knowledge, this is the first PSOCT study to investigate the development of the human brain from infancy to early childhood. Its combination with dMRI further enhances the analysis of fiber tracts and serves as a cross-validation in the developing brain.

### Optical properties revealing myelin development

The myelogenic process begins around birth and continues through adolescence. Our PSOCT study shows substantial development of myelin content in five coronal slabs from 3 to 54 months of age. The optical property of scattering coefficient begins to show a higher signal in the white matter compared to the gray as early as 3 months of age, when myelination is still in its early stages and fibers are largely unmyelinated ^41–43^. Scattering coefficient in the white matter continues to increase over the following 12 months. By 15 months of age the contrast between white and gray matter becomes fully pronounced (Fig. 2). By 4 years of age, scattering coefficient in the white matter increases almost 10 fold (Fig. 3) to approximately 80% of the scattering coefficient reported in the adult human brain ^11,15,44^. The use of optical properties to quantify the myelin content has been validated in previous studies. A quantitative study on PSOCT and histology showed that the scattering coefficient is linearly correlated with the optical density of Gallyas stain, a traditional histological method for myelinated fibers, and this linear relationship holds across multiple brain regions and subjects ^44^.

It is interesting to note that regional differences were also observed in the coronal section at 6 months of age, with higher scattering coefficients in the frontal/parietal lobes than in the temporal lobe. Previous studies have reported that myelin development tends to be from posterior to anterior ^45,46^ and that the occipital lobe is myelinated earlier than the frontal lobe. The present study indicates that the timeline of myelination may also differ between the border of the parietal/frontal lobes (somatosensory/motor cortices) and temporal lobes. Overall, the sensitivity of the optical properties to myelin content suggests that PSOCT may be a useful tool for studying myelination defects in neurodevelopmental disorders.

### Connective pathways in the developing brain

Compared to traditional histology or other forms of microscopy, a distinct advantage of PSOCT is that it has inherent label-free sensitivity to measure the myelin optic axis, allowing for the quantitative determination of fiber orientations across white matter. Anisotropy in the molecular structure of myelin gives rise to birefringence along a unidirectional axis, which is parallel to the axons in the brain. Previous studies using PSOCT or polarized light imaging have shown that the optic axis orientation provides accurate quantification of fiber orientation in the cortex, deep white matter, and subcortical nuclei of the adult human brain ^15,16,47^. In this study, we extended the application of optic axis orientation data in the developing brains with a combined dMRI technique to study the fiber pathways. The optic axis orientation is capable of delineating fiber orientations in infants as young as 3 months of age, despite the low birefringence during the mild myelination stage. Major fiber bundles showed consistency with dMRI tractography for in-plane orientation, during the first 5 years of life. We noticed that noise in the optic axis orientation map is elevated in samples of young age (3 and 6 months old), possibly due to the low birefringence of the white matter. Since the noise in orientation measurements is inversely related to the SNR of the intensity signal ^48^, low birefringence would be expected to lower the SNR of the cross-polarization channel that affects the overall orientation measurements. Nevertheless, the optic axis orientation is a useful tool for studying fiber pathways in the developing brain.

The microscopic resolution of PSOCT reveals complex fiber configurations at finer scales, such as interwoven fibers in the internal capsule and the multiple fiber bands adjacent to it (Fig. 5). The different orientations of these small tracts are appreciated by color-coding the optic axis, which would be lost in the homogenous intensity of scattering otherwise. Tractography applied to the optic axis orientation map differentiates the clusters of fiber bundles running in different directions. One limitation of the current PSOCT technology is that the optic axis depicts only 2D orientation information, which is the projection of the 3D axis into a plane perpendicular to the illumination light. Unlike dMRI tractography that captures the 3D tract orientation, the through-plane angle of fibers is not captured with PSOCT. To map complete connectivity, further advancement of PSOCT technology requires 3D axis orientation measurements, which can be achieved with multiple illumination angles ^49–52^.

### Correlation between PSOCT and dMRI images in brain development

Diffusion MRI (dMRI) provides valuable information in identifying coherent fiber pathways using diffusion properties in the fiber tracts. Using tractography techniques, pathways throughout the white matter of the brain can be depicted in three dimensions ^53,54^. Our study takes advantage of multi-modality techniques to examine the developing brain across scales. Importantly, we use both PSOCT with 10 µm resolution and ex vivo dMRI with 800 µm resolution, spatially co-registered, to investigate the entire coronal section. We found that both optical and diffusion properties reveal developmental patterns that are correlated over the first 5 years of life. However, other than a moderate correlation between PSOCT retardance and dMRI FA, we did not find strong within-subject correlations between PSOCT and dMRI metrics. Correlations between these metrics have been reported in previous studies on adult human brains. The regional variability of quantitative measurements with each modality requires further investigation. Our new technology opens up great possibilities for the study of normal neurodevelopment of the brain and neurodevelopmental disorders.

## Ethics

This study was conducted in accordance with the tenets of the Declaration of Helsinki. The study protocol was approved by Massachusetts General Hospital.

## Data and Code Availability

All software and procedures concerning the data acquisition and analysis have been detailed in the Materials and Methods section. All data sets are available from the corresponding authors upon reasonable request. We will evaluate the request case by case, consulting with the Brain Bank.

## Author Contributions

H.W. (conceptualization, analysis, writing, editing, supervision, funding), N.B. (analysis, writing, editing), D.G. (analysis, writing, editing), A.O. (data collection, analysis, writing, editing, funding), J.L.A.W. (analysis, editing), C.C. (data collection, editing), W.A. (data collection, editing), J.N. (data collection, editing), Z.C. (data collection, editing), A.v.d.K. (data collection, editing), E.T. (conceptualization, analysis, editing, supervision, funding)

## Acknowledgements

This work was supported by the National Institutes of Health (NIH) under award numbers R00EB023993 (H.W.), R21HD106038 (H.W.), U01NS132181 (H.W.), R01NS128843 (H.W.), R01NS109475 (E.T.), R21HD098606 (E.T.) and R21MH118739 (E.T.), and by the MGH Eleanor and Miles Shore Faculty Development Awards Program (H.W.), American Association for Anatomy postdoctoral fellowship (A.O.) and MGH ECOR Interim Support Funding (E.T.).

